# Ultra-high-efficacy Wireless Neuromodulation Using a Conductive Ring Interface

**DOI:** 10.64898/2026.10.19.760169

**Authors:** J. Tian, S. Chen, M. Shotbolt, B. Nandakumar, V. Andre, M. Abdel-Mottaleb, E. Zhu, P. Liang, E. Zhang, P. Ganzer, S. Khizroev

**Affiliations:** College of Engineering, University of Miami, Coral Gables, Florida, U.S.A; Cellular Nanomed, Irvine, California, U.S.A. and University of California, Riverside, CA; Institute of Robotics and Intelligent Systems, Federal Institute of Technology at Zurich (ETHZ), Switzerland; The Miami Project to Cure Paralysis and Department of Biochemistry and Molecular Biology, Miller School of Medicine, University of Miami, Miami, Florida, U.S.A

**Keywords:** brain-computer interface, neuromodulation, neurotechnology

## Abstract

An ab-initio physics study, supported with a COMSOL-based numerical simulation, presents a concept to enable wearable, highly targeted, and ultra-high-efficacy neuromodulation. A target site is defined by a relatively small gap of a miniaturized “ring”-shaped conductive interface implant. If the implant conductivity significantly exceeds the conductivity of the biological tissue, a highly localized electromotive force (emf) is induced in the ring upon application of an alternating current (AC) magnetic field. This emf generates an electric field in the gap, inducing local neural stimulation of the targeted site, e.g., translated to a nerve segment using circumferential cuffs. The wirelessly controlled autonomous stimulator implant acts as a low-resistance nearly-closed path to generate a highly localized electric field on the order of 50 V/m in the gap via application of a 1-kA/m AC magnetic field at a frequency of 10 kHz. The magnetic field is at least three orders of magnitude smaller than the magnetic field required to generate the same electric field strength via conventional transcranial magnetic stimulation. Therefore, this high-resolution, orientation-specific ring neuromodulation system could enable a highly localized, energy-efficient wearable device with significant preclinical and clinical potential.

## 1. Introduction

Neuromodulation by electric fields generated with implanted electrodes is a gold standard in treatment of neurodegenerative diseases and disorders such as Parkinson’s disease (PD), essential tremor (ET), major depression (MD), pain, and others ^1^. Furthermore, implanted neuromodulation is also utilized in several brain-machine interfaces (BMIs) ^2^. Hence, it is difficult to overestimate the significance of any technological advancement in this field. There are several alternative approaches that have been developed to date, however, essentially all of them have run into various stumbling blocks on the way to making neuromodulation technology available to patients and the broader community.

Most commonly, neuromodulation devices use physically implanted wire electrodes to induce electric fields required for stimulation ^3^. In this case, the neuromodulation site is defined by the location of the electrodes. Conductive wires are commonly used to provide the required voltage difference between stimulation electrodes, but this method has the risk of adverse and limiting side effects via off target stimulation ^4^. A less invasive alternative would be the recently emerged percutaneous technology using a conductive polymer material and known as an injectrode. Injectrodes present a safer alternative, minimizing some of the physical risks of the traditional implanted electrodes ^5^. However, because injectrodes still require the use of a bulk conductor for stimulation, they are still prone to some of the same limitations.

Several wireless stimulation techniques have recently emerged for niche applications. These include transcranial magnetic stimulation (TMS) and transcranial direct current stimulation (tDCS) ^6–8^. However, operating in the near-field region, with spatial resolutions on the order of 1 cm, both techniques fundamentally lack the resolution required for substantially more targeted stimulation ^9,10^. TMS, uses rapidly changing (< 100 µs), high-magnitude magnetic fields (>1000 kA/m) to induce eddy currents in relatively large-volume tissue regions (> 1000 cm^3^), with electric fields on the order of 10-100 V/m, as required for exceeding the threshold to stimulate neural tissue. As such, the high magnetic field requirement makes the approach impractical in many potential use cases ^11^. Furthermore, these approaches can also lead to off target effects due to the large volumes of tissues being activated.

Focused ultrasound (FUS) is a promising high-resolution alternative that may be suitable for targeted neuromodulation ^12^. However, FUS suffers from poor penetration depth into the brain and can have side effects due to potential overheating. Other promising, emerging alternatives include optogenetics, magnetothermal, and magnetomechanical stimulation ^13,14^. However, they all are either limited by the need for genetic modification of neural tissue or suffer from low-efficacy energy delivery through skull and/or tissue. Magnetoelectric nanoparticle (MENP) based stimulation, a recently proposed alternative, though promising, still remains in an early phase of development ^15–17^. A millimeter-scale version to use magnetoelectricity as a wireless implant for the endovascular stimulation of a peripheral nerve has also been under consideration, with the energy delivery question remaining open ^18^.

Clearly, there is an unmet need in the field of neuromodulation for an interface that minimizes surgical risk, stimulates neural tissue with a sufficient spatial resolution, and ideally can be activated via a wearable and low-power external device. Using multi-physics COMSOL simulation as a supporting tool, this study aims to exploit electromagnetics physics for creating a technology solution addressing shortcomings of existing alternatives. Specifically, the main goal of the current study is to use a “ring”-shaped conducting interface as a receiving antenna of an applied alternating current (AC) magnetic field ^19^. In this case, the term “ring” refers to any closed (more accurately, near-closed) conducting path with a relatively small gap, as the device can possibly also take the shape of a circle, a triangle, a square, or any other polygon shape, best suited for the considered application. For this work, to simplify the concept description and for the sake of symmetry, only the circular ring shape is discussed. The applied AC magnetic field induces an electromotive force (emf) through the relatively low-resistance closed path in the ring. Then, with the gap length matching the characteristic size of the neuromodulation target (e.g., a segment of the vagus nerve), the electric field due to the emf in the gap can be used as a high-efficacy electric stimulation source. Using this ring system, the eddy currents generated by a traditional TMS approach in a relatively large tissue volume are effectively brought into a highly localized space and along a specific direction determined by the position of the ring and the relative orientation of the gap geometry with respect to the neural tissue being targeted, thus providing significantly improved energy guiding capability, energy transfer efficacy, and spatial resolution. In turn, this capability to convert all the magnetic field flux energy collected by the entire surface area of the ring into the electric field energy in the substantially smaller area defined by the gap allows for significantly reducing the magnetic field strength required for stimulation. The overall purpose of this study is to quantify this ring-enabled electric field localization effect and characterize the overall approach and concept.

## 2. Underlying Physics

### 2.1. Ab Initio Analysis

Before proceeding with the main discussion of the results of a precise COMSOL-based numerical simulation model, we describe the underlying physics via the following ab initio analysis. Similar to conventional TMS, the underlying concept exploits the physics of magnetically induced emf to induce and control the location and direction of eddy currents in conductive biological tissues. In the TMS setup, neural stimulation is achieved by a primary AC electric field induced by an externally applied AC magnetic field along each closed conducting path loop. The emf in each loop (with the resistance 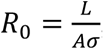, where *L* and *A* are the loop’s length and cross-sectional thickness area, respectively, while *σ* is its conductivity) leads to generation of an electric field, in turn resulting in the eddy current used for stimulation. However, it should be noted that the traditional TMS approach does not allow for the selection of a specific loop required to generate the emf response in the selected target site alone. Instead, an infinite set of closed loops, of comparable conductivity values, are being formed, limited only by the entire conductive body volume (e.g., the tissue around a peripheral nerve). This leads to a set of roadblocks, limiting the capability to target a specific selected stimulation site.

The main hypothesis of this study is to utilize a biocompatible, conductive material to artificially create one specific path for preferentially guiding the eddy current as required for highly targeted stimulation. For example, a conductive polymer could be molded into a ring shape to effectively create a preferred low-resistance closed-loop path. If the conductivity of the closed-loop path is higher than that of the surrounding the neural tissue, the eddy current would be preferentially amplified along this path. Therefore, by placing a target stimulation site along the path, the site would be exposed to the highest possible electric field, with the direction of the stimulation current along the path. Hence, a technical solution would be to open a relatively small physical gap in the artificial closed path, with the gap length large enough to accommodate the targeted neural tissue but small enough to not significantly increase the overall resistance of the “closed” path.

**Figure 1:**
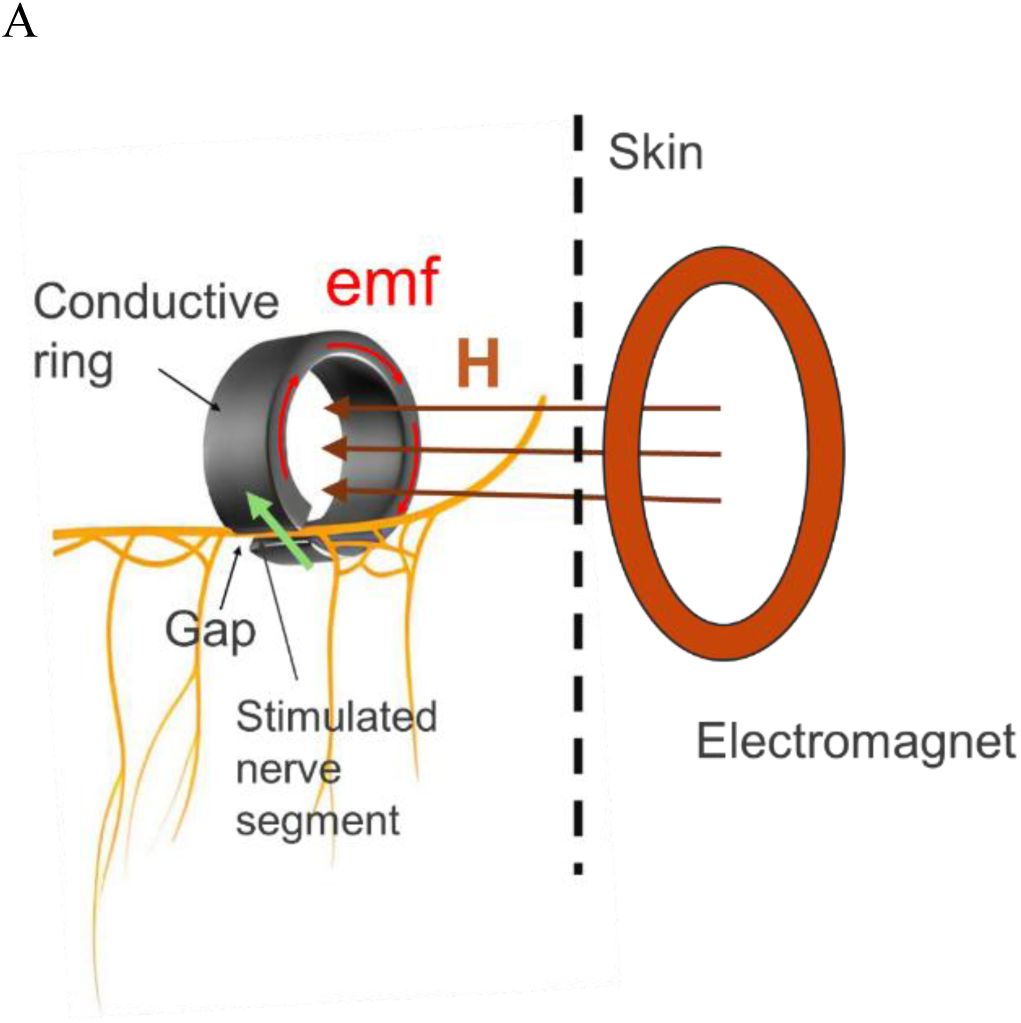
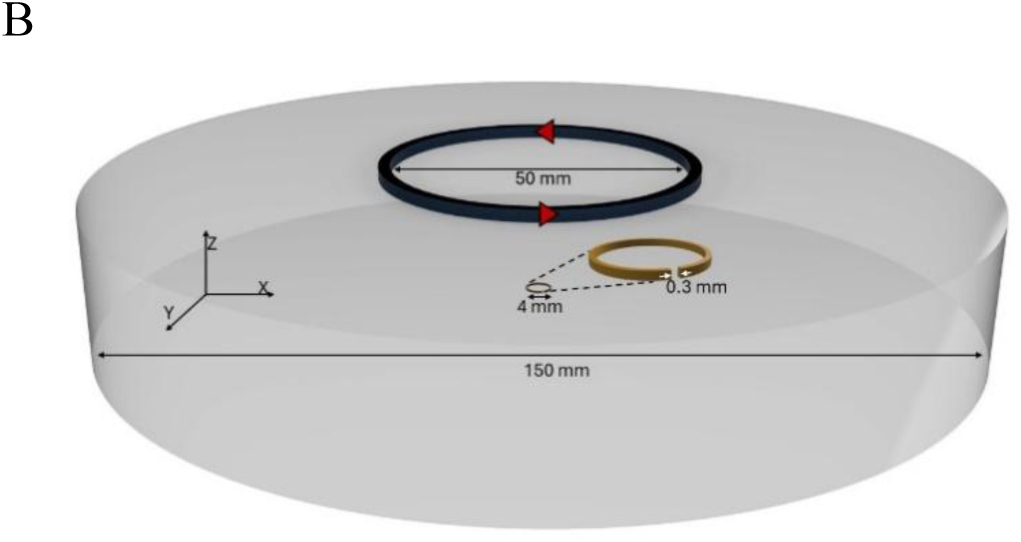
Ring interface. **(A)** High-level illustration of creating a “closed-loop” low-resistance path in the form of a ring, with a small gap to fit the targeted neural tissue for stimulation. The field in the gap, E, is directed along the ring length (green arrow). **(B)** Ring geometry and field orientations used in numerical simulations, for a 4-mm ring external diameter example.

An example of a broken-ring-shaped interface tailored to stimulate a certain region, e.g., the vagus nerve, along a specific direction, is shown in **Figure 1A**. It can be noted that this hypothesis is analogous to the ring-type magnetic head concept used to write information in magnetic recording, with the magnetic flux substituted by the electric current ^20^.

By introducing a relatively small gap, G, in the conductive ring (**Figure 1B**, gold), and assuming there is no leakage current, the modified resistance of the selected loop, R_m_, would be:

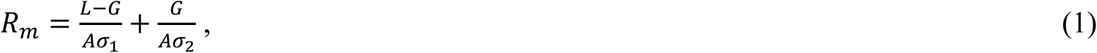

where σ_1_ and σ_2_ are the conductivities of the ring material and the interposed tissue in the gap region, respectively, L and A are the length and cross-section area of the ring wire, respectively. Then, the electric field in the gap, E_G_, could be evaluated via this expression:

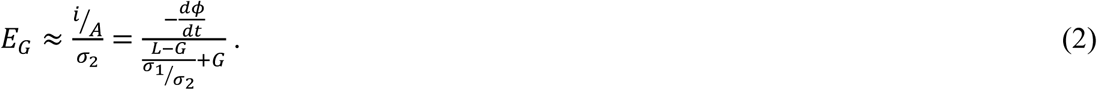

where *i* is the eddy current, *φ* is the magnetic flux through the ring surface. Then, in the limit of σ _1_ / σ _2_ >> 1, the electric field is reduced to 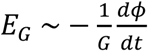.

According to this basic physics analysis, the electric field generated in a 0.3-mm gap of a ring with a radius of 3 mm, upon application of an AC magnetic field with an amplitude of 10 kA/m at a frequency of 10 kHz (equivalent to the TMS pulse rise time of 100 µsec) would be on the order of 50 V/m. This value, 50 V/m, is chosen as a characteristic field intensity needed to stimulate peripheral nerves ^21–24^. For the approximation to be valid, the conductivity of the ring is expected to be at least one order of magnitude higher than the conductivity of the local tissue. Further increasing the radius only by a factor of 3.3, i.e., to approximately 10 mm, would reduce the magnetic field required to induce the required 50-V/m electrical field in the ring’s gap by another order of magnitude, i.e., down to 1 kA/m. The following observation can be made to benchmark this approach against the conventional TMS technique. In the conventional approach, they use a train of short square biphasic pulses (∼100 µs) at a rate ranging from below 0.1 to over 10 pulses/sec, limited by a finite neuron refractory recovery time in the milliseconds range. The pulse rise time is significantly faster than the rate of pulses and can alternatively be described by the rise time, typically in the sub-100-µs range. It can be noted that the rise time ranging from 100 to 10 µs corresponds to the maximum frequency (in the signal frequency spectrum) ranging from 10 to 100 kHz. For clarity, neurons are not receiving 10 kHz trains; instead, they are receiving trains of pulses at regular rates. Reducing the rise time (increasing the maximum frequency) would improve the “squareness” of each pulse. Recent development in the field of TMS, known as Controllable Pulse Parameter TMS (cTMS), showed that reducing the rise time to a value on the order of 10 µs can significantly improve stimulation efficacy and neuronal cell type specificity ^25,26^. Consequently, additionally increasing the maximum frequency to 100 kHz, thus reducing the rise time by a factor of 10, could further decrease the required magnetic field strength. Obviously, this strategy is limited by the inductance of the circuit controlling the electromagnet, the required finite size pulse width (on the order of 100 µs). Coincidentally, the strength and frequency values of this magnetic field harmonic are comparable to those used in modern low-energy portable battery chargers ^27^.

With the main underlying physics described above, the electric fields in the ring’s gap could be further increased using standard incremental methods such as using conductive yokes with non-uniform cross-sections to control current densities, and consequently, electric fields in the gap28.

**Figure 2:**
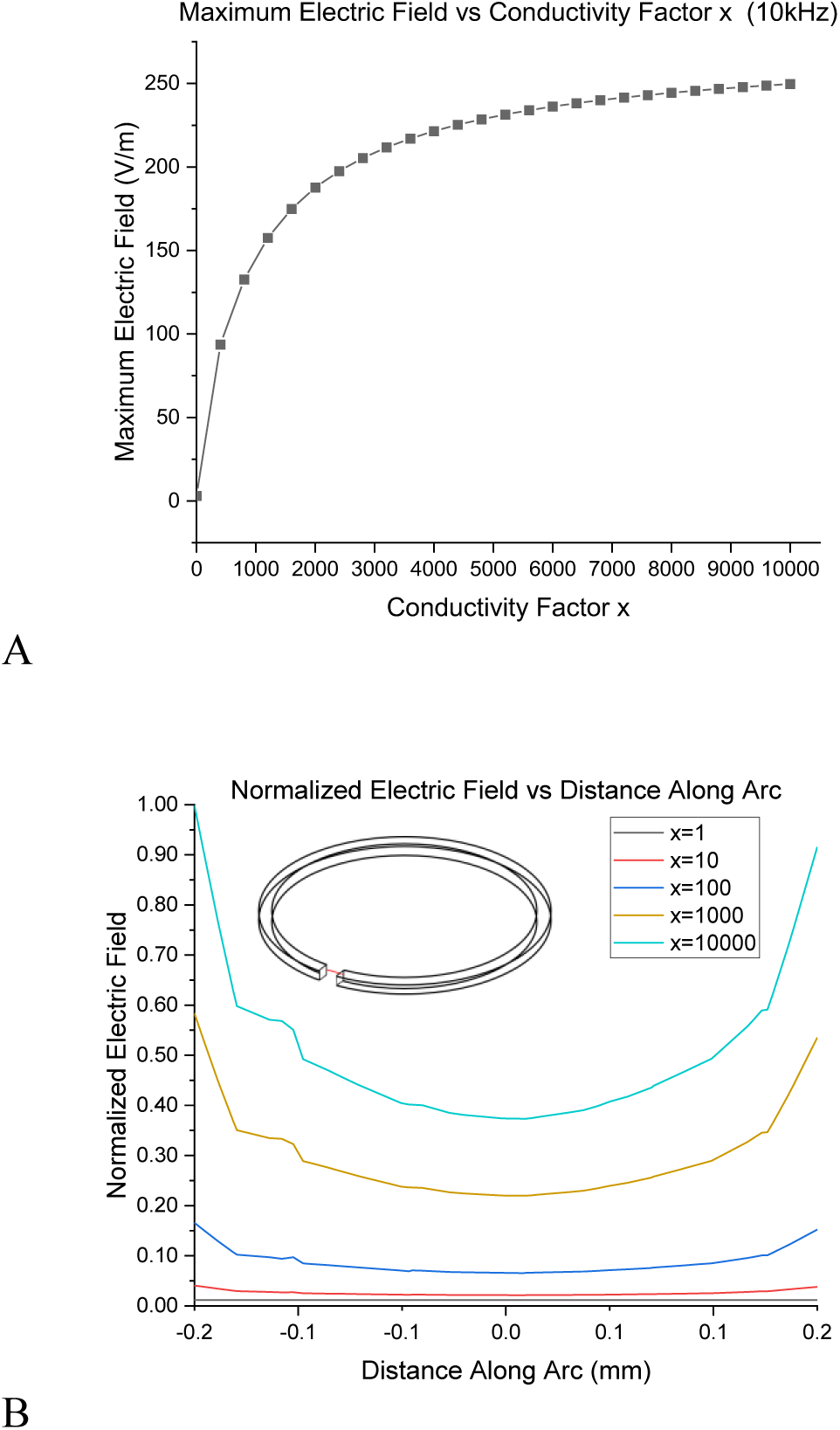
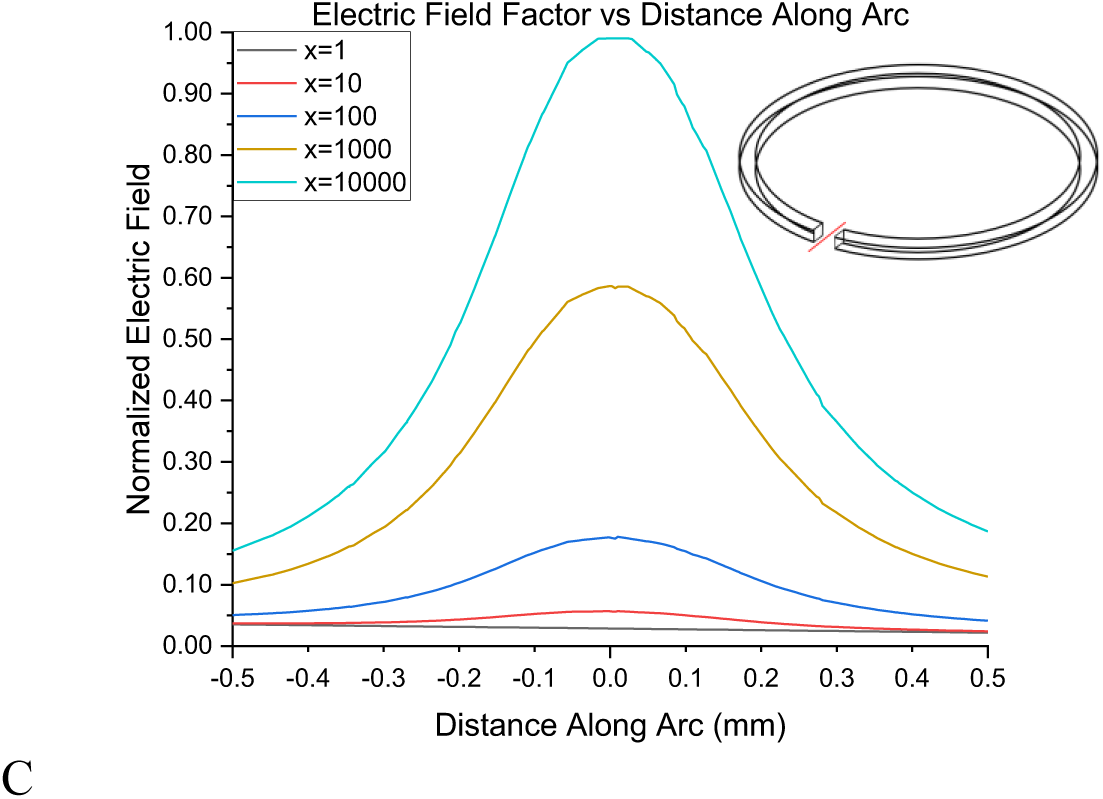
Field in the gap. (**A**) The COMSOL-simulated maximum electric field in the gap as a function of the conductivity factor x=s1/s2 (the ratio of the ring conductivity to the average tissue conductivity). The applied AC magnetic field strength at 10 kHz is approximately 16 kA/m. (**B**). The normalized electric field along the gap length versus the distance along a central line along the Y-axis (red reference line within the gap). (**C**). The normalized electric field along the gap length versus the distance along a central line along the X-axis, between the gap edges (red reference line within the gap).

### 2.1. COMSOL Simulations

The discussed numerical simulation model is designed to more precisely validate our overall hypothesis, compared to the above ab initio analysis. For example, the ab initio analysis does not consider capacitive effects due to dielectric properties in complex geometries, nor does it consider inductive effects due to non-circular shapes. As a result, the COMSOL simulations are vital to provide the frequency response and resulting non-linearities of real-life systems. Given the circular symmetry, a trivial circular coil electromagnet was chosen to generate a magnetic field perpendicular to the main surface of the ring neuromodulation interface (**Figure 1B**). The coil diameter is selected to be substantially larger than both the outer diameter of the ring and the distance between the ring and the electromagnet. For example, if both the ring diameter and the distance between the ring and the electromagnet are on the same order of magnitude, e.g., approximately 10 mm, the coil would need to be at least 20 mm in diameter. This ensures the magnetic field perpendicular to the ring does not significantly vary over the surface of the ring, thus making the design more predictable. Here, it is assumed that the electromagnet and the ring are co-axially aligned, as shown in **Figure 1B**, though providing precise alignment is not critical, according to the underlying physics. Throughout the simulation, the ring gap length, G, was kept constant at 0.3 mm, i.e., comparable to the characteristic size of a theoretical nerve stimulation target. As explained above, the frequency bandwidth under study, 0-100 kHz, was chosen to provide an adequate fit of the biphasic signal waveform used in conventional TMS, given the most recent development, known as cTMS, according to which reducing the rise time to a value on the order of 10 µs was shown to improve the stimulation efficacy and cell-type specificity ^26^.

For the initial testing, the electric field in the gap was calculated as a function of the ratio of the ring conductivity σ_1_ and the average neural tissue conductivity σ_2_, x = σ_1_/σ_2_, further referred to as the conductivity factor x. The gap cross-section was chosen to be square, with a side of G*k, and a default value of k=2/3, where k is ratio of the square side to the gap length G. The results are summarized in **Figure 2**. As expected according to the above physics considerations (Equation 2), the electric field increases with the ratio x. It can be noted that increasing the ring conductivity by three orders of magnitude compared to that of the neural tissue should provide a notable effect.

**Figure 3:**
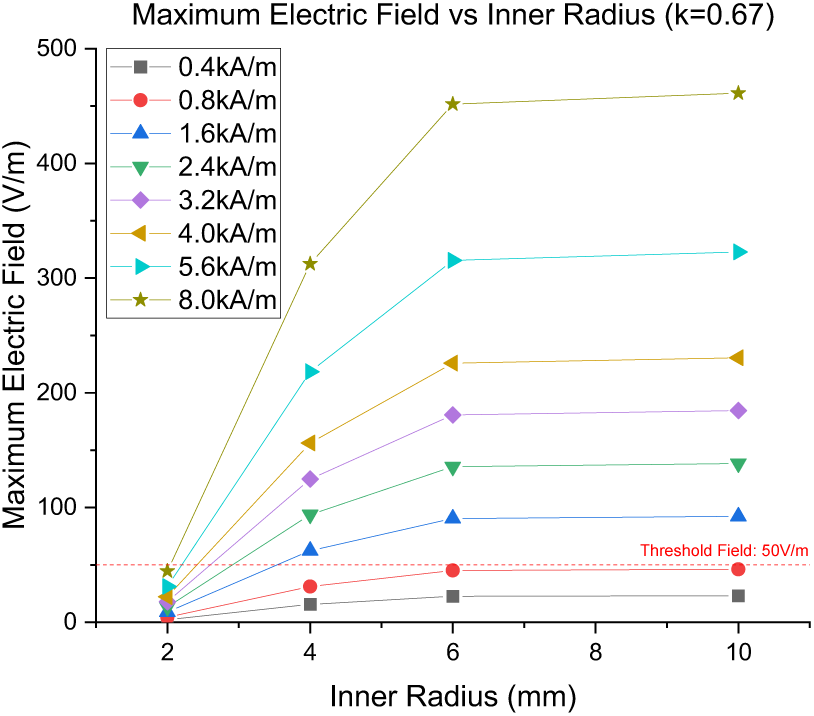
Field versus gap radius. The COMSOL-simulated maximum electric field in the gap as a function of the ring’s inner radius (mm) across a range of magnetic field strengths. The AC magnetic field frequency applied to the ring is 10 kHz (threshold field = induced electrical field needed for neural tissue stimulation). k=2/3 (the gap square cross-section side is 0.2 mm)

Next, the electric field in the gap was calculated as a function of the ring’s inner radius, with k held constant at 2/3 (∼0.67). The results are shown in **Figure 3**. It can be noted that the field starts to saturate after the radius is increased above 10 mm. This trend is in agreement with the above physics consideration (Equation 3), that the modified resistance of the loop increases with the inner radius at a higher rate when the cross-sectional area, A, is smaller. Another contribution to the observed non-linearity is the fact that the inductance of the path increases with the radius increase, thus effectively further increasing the high-frequency impedance.

The electric field in the gap was then evaluated as a function of k, with the inner radius held constant at 2 mm (**Figure 4**). The induced field strength does not saturate after k is increased above approximately 6, indicating there is room for further optimization. This trend agrees with the above physics consideration (Equation 2), assuming lossless energy transfer in the near-closed loop (i.e., the ring). Indeed, the larger the cross-sectional area of the ring, the more justified the lossless current flow approximation is. In the lossless flow regime, the field does not depend on the cross-section, provided the cross-section is uniform along the entire closed path. In a more realistic case, when the cross-section is relatively narrow, compared to the gap length, the path becomes more open, thus leading to increased current losses and smaller electric field in the gap.

The electric field in the gap for the ring parameters, optimized through the above calculations for the maximum electric field in the gap, was calculated as a function of the perpendicular magnetic field in the center of the ring (**Figure. 5**).

**Figure 4:**
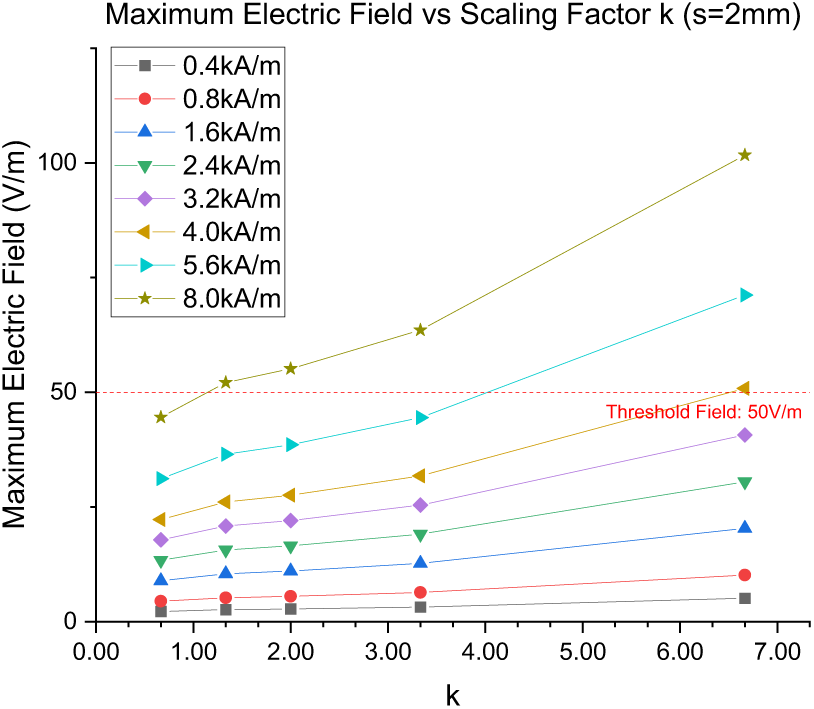
Field versus scaling factor. The COMSOL-simulated maximum electric field in the gap as a function of the scaling factor k. The AC frequency is 10 kHz. The inner radius, s, is 2 mm.

**Figure 5:**
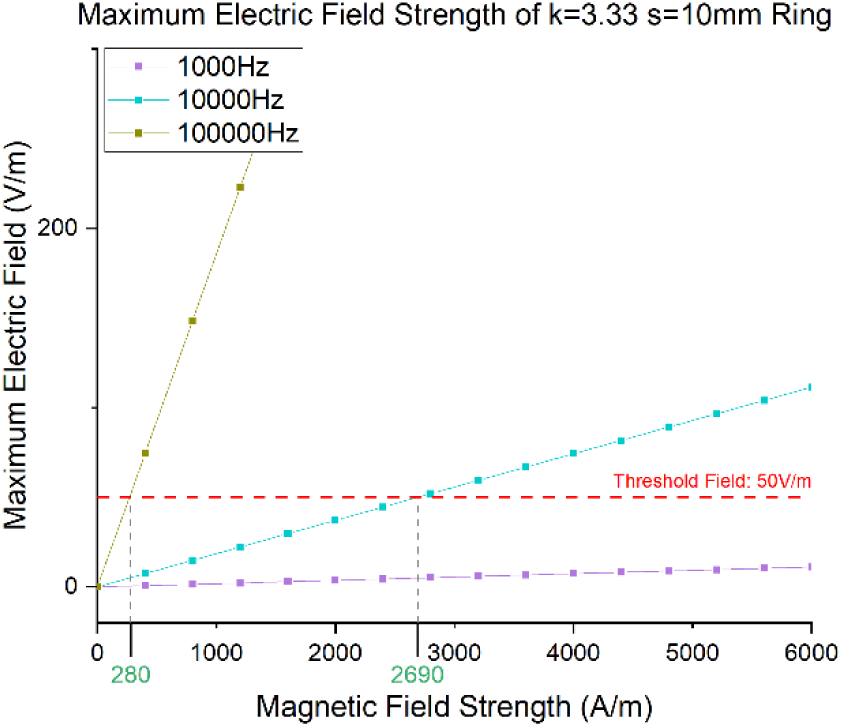
Electric field in the gap verus magnetic field. The COMSOL-simulated maximum electric field in the gap as a function of the perpendicular magnetic field in the center of the ring. k=10/3 (the gap square cross-section side = 1 mm). s=10 mm.

**TABLE 1.** Comparison of the studied ring interface and the state-of-the-art TMS.

|  | Ring Interface | TMS |
| --- | --- | --- |
| Source H (T) required for inducing E of 50 V/m | ~0.1 kA/m (can operate at a significant frequency of 100 kHz) | ~1000 kA/m (at 10 usec pulse widths) |
| Stimulation region size | ~ 0.1 mm | ~ 100 mm |
| Stimulation orientation | Well defined (perpendicular to the ring plane) multiple rings can potentially enable spatially selective modulation | Not defined |
| Power source dimensions | Hand-held battery-sourced, cost-effective, can be wearable | Bulky and requires capacitor banks |
| Power requirements | Low (mWs -Ws) (identical to the power consumption of wireless energy chargers) | High (1000s W) |
| Working frequency | Dynamic coupling frequency dependent on the ring's geometry from 10 kHz to 100 kHz | 10 kHz (ramp rate: 100 μs) (can range from below 50 to above 100 μs) |
| Stimulation pulses | Continuous stimulation is achievable | Sporadic pulses limited by the charging sequence of the capacitor bank |
| Invasiveness | Low to Moderate (implant is used) | None (no implant is required) |

According to the simulation results, a magnetic field on the order of 2.7 kA/m at 10 kHz would lead to an electric current in the gap of the ring on the order of 50 V/m, which is roughly the threshold required to stimulate a representative peripheral nerve ^21,24^ (Figure 5, cyan line).

**Figure 6:**
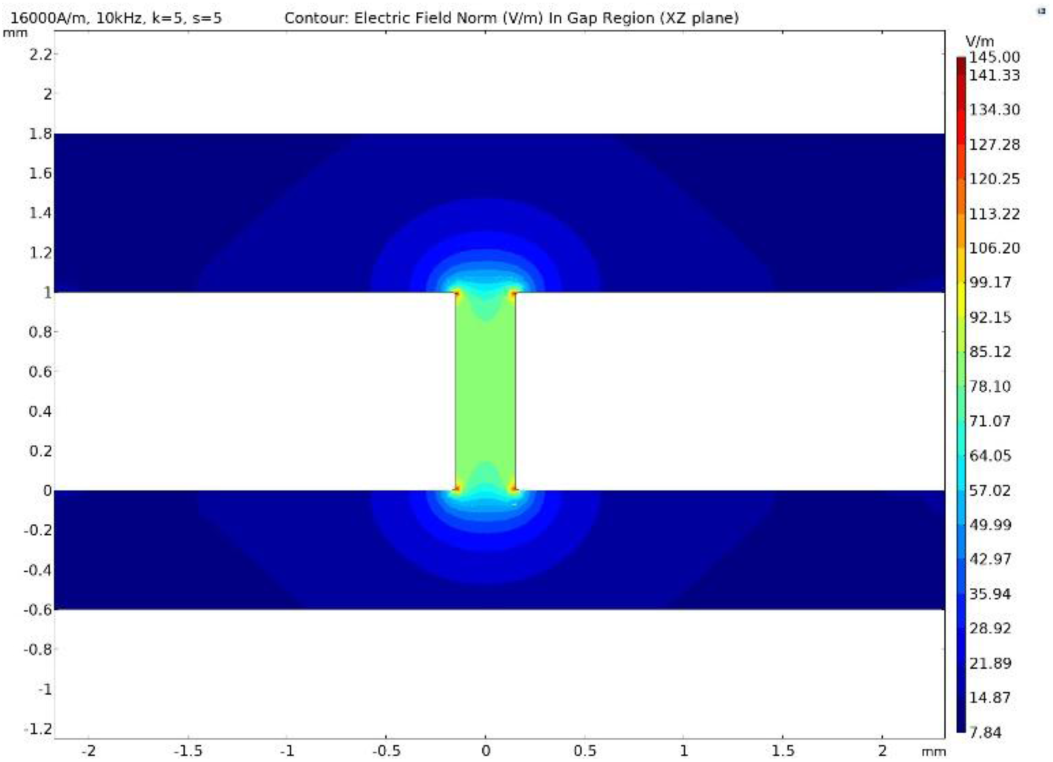
Contour map of the electric field in the gap. The contour map of the electric field strength in the gap, view is normal to the XZ plane. Note that the electric field in the gap is above 50 V/m within, while hotspots exist at the corners.

**TABLE 2.** Electrical Properties of materials and meshing for simulation.

| Material | Electrical Conductivity (S/m) | Relative Permittivity | Meshing | Meshing Preset | Minimum Element Size |
| --- | --- | --- | --- | --- | --- |
| Ring Material | Variant | 1 | Mesh 1 | Semiconductor (extremely fine) | 70μm |
| Tissue (phantom) | 1 | 1 | Mesh 2 | Semiconductor (extra fine) | 500μm |
|  |  |  | Mesh 3 | Custom | 2mm |
| Air | 1E-9 | 1 | Mesh 4 | Custom | 5mm |

The main theoretical result that such low-field, high-frequency, magnetic field pulses can indeed induce suprathreshold firing was confirmed via basic in vivo experiments in Wistar rats (**Supporting Material Fig. S1 and three videos VidS1, VidS2, and VidS3**).

## 3. Methods

### 3.1. Simulations

The AC/DC - Magnetic Fields (mf) module from COMSOL was used to perform magnetic field simulation with low frequency. The physical components of the basic simulation model consist of three parts: the magnetic field source, the conductive ring, and the surrounding tissue. The AC Magnetic Field Source: A frequency domain study to simulate dynamic effects was used following a stationary study to solve for the initial solution of each specific field parameter set. As a benchmark set of field specifications, conventional TMS waveforms specs were used. For example, a typical biphasic TMS signal waveform has a field rise time of 100 µs, corresponding to the significant value of 10 kHz in the signal frequency spectrum. Hence, by default the dynamic response was studied at the maximum frequency of 10 kHz. Furthermore, given that the current approach allows to substantially lower the applied magnetic field strength, thus allowing to reduce the inductance of the system and, consequently, increase the field frequency, compared to the conventional TMS, with the goal to further improve the biphasic waveform generation, a ten-fold increase of the maximum frequency, i.e., to 100 kHz, corresponding to the rise time of 10 µs, was also considered as a case study. As discussed above, this time rise corresponds to the recent development in the TMS area known as cTMS. The magnetic field source was modeled using two different methods: a physically modelled coil using homogenized multiturn coil (numerical) formulation with properly calculated current with input current to generate the desired magnetic field magnitude at the center of the ring gap. While this method is physically realistic and can capture TSM behavior more accurately, it does significantly increase computational load and introduces field artifacts at air–tissue interfaces in close proximity of the source coil. The second approach used Dirichlet boundary conditions to impose a uniform magnetic field across the entire domain. In this configuration, a cylindrical air domain was used to enclose the simulation region, and the H_z_ component (in A/m) was directly specified on the side boundary. Using Dirichlet H field condition with a cylindrical air domain, which shares circular symmetry with the ring structure, will also prevent corner artifacts otherwise introduced with rectangular or cubical domains. The Dirichlet condition is particularly effective for frequency-domain and field strength sweep simulations, offering a simplified yet accurate approximation of a uniform far-field magnetic excitation. After testing both methods under identical conditions for the initial conductivity ratio study, we observed only minor differences (<1%) in the induced electric field within the ring gap, which is the region of interest. Therefore, the computationally less burdensome Dirichlet-based method was adopted for all subsequent simulations.

The Conductive Ring: The geometry of the conductive ring directly determines the electric field generated by the induced emf, as does the material property of the ring. In this scenario, the region of interest is within the gap of the ring, where maximum induced electric field is expected to occur. The ring material has electrical conductivity of σ_1_=xσ_2_, with x being the dimensionless conductivity factor, indicating the difference compared to the characteristic biological tissue conductivity on the order of 1 S/m. For example, an x value of 1000 would lead to a ring conductivity of 1000 S/m, which would fall into the currently or potentially achievable range of typical biocompatible conductive polymers like Polyaniline (PANI), Polypyrrole (Ppy) or the widely used poly(3,4-ethylenedioxythiophene) polystyrene sulfonate (PEDOT:PSS), heavily depending on the doping and processing methods ^29^. It is worthwhile mentioning that PEDOT:PSS has demonstrated conductivity on this order of magnitude in flexible and stretchable bioelectronic interfaces ^30–32^. The ring gap is set to 300 µm to match the approximate cross-sectional size of a targeted peripheral nerve bundle.

The Surrounding Tissue: At this early stage of the development, the surrounding tissue was simulated as a homogeneous phantom, with values for the averaged conductivity and permittivity of approximately 1 S/m (σ2) and 3.5x10^4^, respectively ^33,34^. Additionally, computations with human tissue models are subject to singularities existing in the form of artifacts, with the occurrence most frequently at the interfaces between regions of drastically differing conductivity. The homogenous model was chosen to ensure the anticipated local maxima will be guaranteed at ring/tissue interface and not be overridden by the described numerical artifact (as shown in Figure 6) ^35^. To help with numerical convergence and avoid singularity related issues in the solver, the conductivity of the surrounding air domain was set to a small but non-zero value of 1 E-9 S/m, which is a common approximation in low-frequency electromagnetic simulations using Magnetic Fields (mf) module.

The meshing setting follows a 4-level hierarchy. The conductive ring and the immediate surrounding tissue, including the gap region of interest, was meshed with semiconductor settings preset to “extremely fine”, with minimum element size of 70um across the 300µm gap to ensure field smoothness, this mesh has the highest meshing priority (mesh 1). Due to the lack of symmetry in the geometry, free tetrahedral meshing was used. The surrounding homogeneous tissue domain to air domain at boundary of simulation was divided into three additional regions (mesh 2, mesh 3, and mesh 4) with progressively larger element sizes as the distance from the ring gap increases. To maintain smooth transitions and avoid abrupt changes in element size (which may impair solver stability), the mesh growth factors were set to 1.05, 1.08, 1.20, and 1.20, respectively. The meshing strategy enabled accurate simulation of the localized E-field within the region of interest (ring gap) without compromising convergence or introducing expensive computational workload.

All simulations were performed using a frequency-domain study with the BiCGStab iterative solver, configured with a relative tolerance of 0.001. This setting was used for the quasi-static regime under investigation (10–100 kHz). Within the homogeneous domain, no field singularities or unexpected local maxima/minima were observed. The induced electric field varied smoothly across the ring gap, and the direction and magnitude of the field remained consistent with theoretical expectations. The small element size of 70 µm across a 300 µm gap further ensured adequate spatial resolution in the critical field concentration zone.

## 4. Discussion

Both the presented ab-initio physics analysis and the COMSOL-based numerical simulations support the hypothesis that the ring interface can induce electric fields required for suprathreshold activation, localized in the target site with the characteristic size defined by the ring’s gap length. Although, to simplify the concept description, the simulation focused on the gap value of 0.3 mm, tailored to match a targeted segment of a hypothetical peripheral nerve, it could be extended to a wide range of gap lengths, specific to a selected target site. It is also possible that in the future this approach could be translated to target sites in the central nervous system. Not requiring implanting any active electronic devices, e.g., transistors for energy harvesting, the approach is relatively straightforward to implement surgically. The simulation confirmed the theoretical prediction that application of a relatively weak AC magnetic field can generate a relatively high eddy current in the gap, exceeding the threshold required for local induction of action potentials. Notably, this statement was also confirmed via preliminary in vivo experiments on rats (**Supporting Material**). Noteworthy, the conventional TMS system would require at least three orders of magnitude stronger magnetic field to induce a comparable electric field strength and thus a comparable stimulation current. As also predicted through the ab initio physics calculations, the significant boost in the device effectiveness is due to the fact that the ring interface is designed to convert all the magnetic energy, collected from the AC magnetic flux through the ring surface area in the near-field region, into an induced electric field localized into the significantly smaller area defined by the ring’s cross-sectional area and gap, with the magnetic-to-electric-energy conversion accomplished via the emf in the near-closed path defined by the ring. The physics analysis also showed that for a ring material with a relatively high conductivity value, on the order of 10^5^ S/m, the electric field in the gap, induced via application of a relatively small magnetic field on the order of 0.5 kA/m with the rise time of 10 µs, can surpass neural stimulation threshold by a large margin. Another important advantage of the ring interface approach, besides the great efficiency boost, is the prospect of using a small (radius) electromagnetic coil to generate the relatively small required magnetic field, even at a physiologically relevant distance away from the coil. In turn, the lower inductance of the small coil makes it easier to use a larger maximum frequency, thus improving the rise time, as required for generating sufficiently high electric currents localized to a gap-long site. Overall, due to the combination of this boost in the field conversion efficiency and the ease to operate at significantly higher frequencies, the ring interface approach could be implemented with a significantly reduced form factor, with power specs and frequency characteristics matched to those used in the conventional low-power wireless portable energy chargers, thus potentially unlocking a new realm of wearable medical device applications. The stimulation depth is limited by the trade-off between the inductance of the circuit used to control the electromagnetic coil and the radius of the electromagnet. The larger the field penetration depth, the larger the required electromagnet coil radius. However, the larger the radius, the larger the inductance and thus the smaller the maximum frequency. For example, the magnetic field on the order of 1 kA/m at a maximum frequency of 100 kHz, as required to induce the stimulation threshold electric field in the gap according to the theory, could be generated by a 10-cm electromagnet with a 1000 A·turn current approximately 10 cm away from the coil.

Given the high-frequency operation of the medical device, for the analysis to be valid, the thickness of the ring should not exceed the skin depth. Indeed, for the conductivity value of 10000 S/m, the skin depth at the frequency of 100 kHz is on the order of 7 mm, thus meeting the requirement.

Furthermore, the following conservative thermal management analysis can be used to show that in this miniaturized implementation, under equilibrium conditions, the implant’s temperature increase would not exceed even a millionth of a degree, thus being lower than the thermal noise in the system. Indeed, in the worst-case scenario estimate, it can be assumed that the entire power delivered by the AC magnetic field is given to the ring and dissipated into heat (Joule’s heating), while no sink is used to remove the heat, 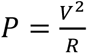, where V is the induced emf and R is the electrical resistance of the ring ^36^. Then, under equilibrium, the ring’s temperature increase is 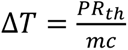, where m is the mass of the ring, R_th_ is the thermal resistance ^37^. Assuming R_th_ = d/K, where K is the thermal conductivity and d is the ring thickness, 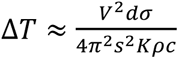. For example, for PEDOT:PSS (s ∼ 1 x 10^5^ S/m, K ∼ 1 W m^-1^ K^-1^, C_P_ ∼ 1.5 x 10^3^ J kg^-1^ K^-1^, r ∼ 1.6 x 10^3^ kg m^-3^, for the ring with s = 5 x 10^-3^ m and d = G= 3 x 10^-4^ m, assuming that the emf can induce an electric field in the gap close to the neural stimulation threshold value of approximately 50 V/m, the voltage drop in the gap would be V = 1.5 x 10^-3^ V. Then, according to this equiation, the temperature of the ring will go up only by a millionth of a kelvin, which is a negligible amount.

In summary, this novel physics-based ring concept paves the way for enabling an extremely miniaturized and energy-efficient, injectable and with no active components, wireless interface, coupled with a wearable external device that can generate highly targeted neuromodulation with stimulation levels on par with or better than those generated by a state-of-the-art TMS system (**Table 1**).

Current peripheral neural stimulation interfaces have been efficacious in treating a number of diseases but still suffer from an array of shortcomings ^38^. In theory, it is possible that the ring interface assessed here can be placed using a percutaneous approach, therefore mitigating the risks involved with a more invasive surgery ^39,40^. In addition, several current peripheral nerve stimulation interfaces are also negatively affected by lead migration ^41^. The proposed ring interface, although seemingly leadless, may still possibly undergo some degree of migration without specific anchoring (especially in an area of the body that is exposed to flexion, extension, strain, and / or rotation that can lead to temporary and reversible migration as the surrounding tissue is affected). Regardless, the ring interface studied here is not tethered to an implanted pulse generator and therefore should exhibit less risk of large-scale migration over time. The gap size of the ring can likely take on a wide range of values, given the selected aspects of the overall interface components. This can have a significant effect on what tissues can be reasonably targeted for stimulation. In addition, future studies will need to assess what degree of device encapsulation would occur using different materials for the ring, as this can have an important effect on the parameters required to enable neural stimulation using the ring interface. Importantly, the dramatically enhanced spatial resolution of neural stimulation at dramatically reduced applied magnetic field amplitudes, when compared to TMS, represents a promising feature of the proposed interface. These and other critical factors await demonstration *in vivo* to validate the modeling predictions outlined here to further demonstrate feasibility. Furthermore, it is noteworthy that, in the future, the described ring interface technology could be scaled into a wirelessly controlled network of many (thousands or more) sub-millimeter size autonomous stimulators, in one implementation being randomly scattered into a target region or, in another implementation, tethered to a specific location. According to a direct scaling principle, a millimeter-diameter ring with a 100-µm thickness would require a sub-100-µm gap to locally induce electric fields sufficient for stimulation under the above applied magnetic field conditions. Such dimensions can be achieved using conventional printing/lithographic techniques ^42^.

In summary, by providing highly localized neuromodulation at orders-of-magnitude lower power than conventional TMS without auxiliary circuitry, this system offers a promising pathway toward wearable preclinical and clinical applications.

## Supporting information

S1: 22kHz, sinusoidal, 500 cycles, Vpp=3.1V

S2: 22kHz, sinusoidal, 500 cycles, Vpp=3.25V

S3: 22kHz, sinusoidal, 500 cycles, Vpp=6.0V

## 6. Conflicts of Interest

Authors declare no conflicts of interest.

## 7. Author Contributions

JT conducted numerical simulations with COMSOL, prepared figures, designed amplifier used in the reduction of practice experiment, and helped with writing the manuscript. SC helped build geometrically complex parts of the numerical model using a 3D CAD software package SOLIDWORKS. MA, MS helped with COMSOL simulations, preparation of figures, data analysis of experiments. BN helped with preparation of figures and design of experiments, conducted reduction-to-practice experiments. EZhang, EZhu, VA, and PL helped with writing the manuscript. PG co-invented the concept, helped design the experiments and write the manuscript. SK oversaw the entire project, invented the concept, designed the experiments, and wrote the paper.

## Acknowledgements

This study was partially supported through National Science Foundation (NSF) under grant number ECCS-211082, and the Sylvester Comprehensive Cancer Center (SCCC) intramural funding program under the core National Institutes of Health (NIH) grant 5P30240139-02.

## Data Availability Statement

Available upon request.

## SUPPORTING MATERIAL

### S1. Reduction to Practice: In Vivo Experiments Showing Three Orders of Magnitude Less Field (Compared to TMS) Required to Induce Suprathreshold Activation

To translate the proposed concept into practice at this early stage of development, we conducted proof-of-principle experiments in Wistar rats (University of Miami IACUC#21-175). The objective was to verify if the electric field generated within the gap of the ring structure was in the right order of magnitude range to induce suprathreshold neural activation using an AC magnetic field with an amplitude orders of magnitude smaller than that employed in conventional transcranial magnetic stimulation (TMS) systems (∼1000 kA/m).

**Figure S1:**
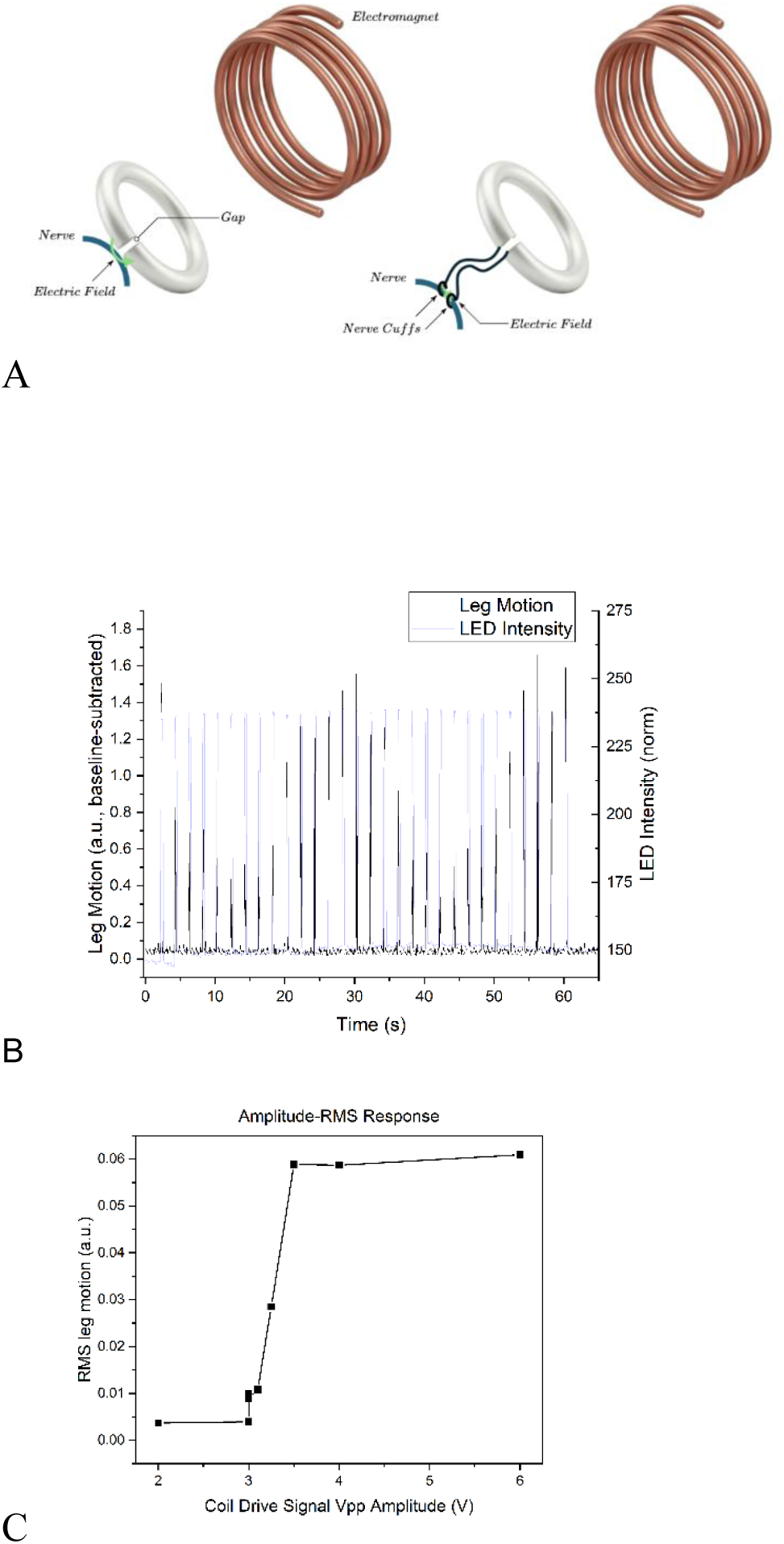
In vivo experimental validation and reduction to practice. (**A**) Comparison between the computational model of a ring receiver directly coupled to a nerve segment (left) and the *in vivo* experimental setup (right). In the experimental configuration, a proof-of-concept monolithic metallic ring interfaces with the nerve via a pair of circumferential cuffs, with the gap impedance matched to the simulated ring gap to account for biological tissue as load. (**B**) Recorded leg motion response (black) synchronized with stimulus onset (light blue) during a stimulation session (Vpp = 6V, f = 22 kHz, sinusoidal). (**C**) Muscle movement showing the RMS leg motion amplitude as a function of the coil drive signal amplitude (Vpp), demonstrating the threshold and saturation characteristics of nerve response.

To evaluate the feasibility the proposed concept, we conducted proof-of-principle experiments in Wistar rats (University of Miami IACUC#21-175) (**Figure S1**). The objective was to verify if the electric field generated within the gap of the ring structure was in the right order of magnitude range to induce suprathreshold neural activation using an AC magnetic field with an amplitude orders of magnitude smaller than that employed in conventional transcranial magnetic stimulation (TMS) systems (∼1000 kA/m). Prior to magnetic field stimulation, nerve engagement was quantified using conventional electrical stimulation. A train of 20 Hz pulses (200 μs pulse width, 0.5–5 mA) was delivered through a nerve cuff using an isolated constant-current stimulator (AM-Systems). This resulted in reliable target engagement of the gastrocnemius–soleus and anterior tibial muscles, producing ankle plantar flexion. Notably, currents as low as 0.5 mA were sufficient to induce isolated flexion of the digits in the hindlimb. This electrical stimulation protocol served to confirm proper operation and effective engagement of the nerve-cuff interface. Following confirmation of electrical activation, we proceeded with experiments using a high-frequency, low-power electromagnetic stimulator. The available low-pass-filtered power source for magnetic stimulation could generate a maximum magnetic field strength of only ∼0.5 kA/m at 10 kHz. However, numerical simulations indicated that, to induce an electric field exceeding the estimated neural activation threshold of ∼50 V/m within the gap, an AC magnetic field on the order of ∼5 kA/m would be required. To satisfy this modeling-derived requirement with an appropriate safety margin, we modified the coil geometry by increasing the number of turns from one to three and enlarging the ring radius by a factor of two. With the primary goal of demonstrating that sufficient energy could be delivered to a relatively short nerve segment (∼1 mm) under these conditions, and with biological tissue present in the gap, we connected low-impedance copper leads from the sides of the gap to a tissue segment using two nerve cuffs separated by approximately 1 mm (**Figure S1A**). In this configuration, the majority of the impedance was concentrated within the biotissue (nerve) region, thereby satisfying the modeled boundary conditions. Representative experimental results obtained from three animals are summarized in **Figure S1B**. The vertical axis shows limb displacement in response to application of the magnetic field. A clearly discernible, isolated flexion of the hindlimb digits was observed, strongly time-correlated with magnetic field application and comparable in magnitude to the response elicited by electrode-based electrical stimulation. The root-mean-square (RMS) displacement amplitude as a function of the electromagnetic coil drive peak-to-peak voltage is presented in **Figure S1C**. Overall, these results supported the theoretical prediction that high-frequency (>10 kHz) magnetic fields with amplitudes three orders of magnitude lower than those used in conventional TMS, corresponding to magnetic field rise times on the order of 100 μs, can achieve reliable, localized neural activation.

#### Videos of In Vivo Experiments

Three videos of the in vivo functional are included in the supplemental materials. A conductive ring was connected to the sciatic nerve of a Wistar rat through nerve cuff. The stimulus consists of a 22 kHz sine wave pulse train (500 cycles) at three coil drive voltage amplitudes, 3.1, 3.25 and 6 V, in video files **VidS1**, **VidS2** and **VidS3**, respectively. Clear, observable isolated flexion of the hindlimb digits are synchronized with the onset of the pulse train (indicated by the green LED).

### Methods

#### In vivo experiments

In this experiment, adult male Wistar rats weighing 300-400 grams were anesthetized with urethane (1.6g/kg i.p) and implanted with a nerve cuff around the sciatic nerve of the hindlimb. First to quantify nerve engagement, electrical stimulation (a train of 20 Hz/200 microsecond pulses of a 0.5-5mA current) was delivered to the nerve cuff using an isolated constant current stimulator (AM-systems), resulting in effective target engagement involving gastrocnemius-soleus and anterior tibial muscle causing ankle plantar flexion. A current of 0.5mA enabled isolated flexion of digits in the hindlimb. Once the electrical activation was used to confirm the operation and engagement of nerve cuff interface, we conducted experiments using the high frequency low power electromagnetic stimulator.

The induction coil in the reduction to practice in vivo experiment was driven by a custom-designed high-power Class AB amplifier using a PA09 monolithic operational amplifier (Apex Microtechnology), selected for its high slew rate and ability to drive reactive loads. To ensure stability and precision when driving the inductive air-core electromagnetic coil, the system was configured as a voltage-to-current transconductance setup (220mA/V, flatband). Prior to *in vivo* application, characterization was performed to establish the transconductance gain and frequency response of the assembly. Given the Hall-effect sensors were limited by finite sensing slew rates and could not accurately resolve the high-frequency magnetic fields used in this study, the generated magnetic field was estimated analytically using quasi-static assumption, the instantaneous magnetic field was calculated using the Biot-Savart law based on the instantaneous current measured across a precision non-inductive sensing element in series with the air core coil. This approach provided a reliable estimate of the field intensity and coupling potential during the pulse train onsets observed in the experimental video.

## Notes

### Competing Interest Statement

The authors have declared no competing interest.

